# Temporal and spatial niche partitioning in a retrotransposon community of the Drosophila genome

**DOI:** 10.1101/2024.08.14.607943

**Authors:** Marion Varoqui, Mourdas Mohamed, Bruno Mugat, Daniel Gourion, Maëlys Lemoine, Alain Pélisson, Charlotte Grimaud, Séverine Chambeyron

## Abstract

Transposable elements (TEs), widespread genetic parasites, pose potential threats to the stability of their host genomes. Hence, the interactions observed today between TEs and their host genomes, as well as among the different TE species coexisting in the same host, likely reflect those that did not lead to the extinction of either the host or the TEs. It is not clear to what extent the expression and integration steps of the TE replication cycles are involved in this ‘peaceful’ coexistence. Here, we show that four Drosophila LTR RetroTransposable Elements (LTR-RTEs), although sharing the same overall integration mechanism, preferentially integrate into distinct open chromatin domains of the host germline. Notably, the differential expressions of the gtwin and ZAM LTR-RTEs in ovarian and embryonic somatic tissues, respectively, result in differential integration timings and targeting of accessible chromatin landscapes that differ between early and late embryonic nuclei, highlighting connections between temporal and spatial LTR-RTEs niche partitionings.

## Introduction

Proper development of multicellular organisms relies on the temporally and spatially regulated expression of genes encoded by the genome. However, not every DNA sequence, even if it can be expressed within a genome, plays an indispensable role. Some sequences, such as transposable elements (TEs) exhibit a self-serving behavior due to their ability to autonomously express and insert in various locations within the genome and can be considered as genomic parasites. Indeed, continuous TE activity may lead to harmful mutations, including disruptions of coding sequences, chromosomal rearrangements facilitated by ectopic recombination, and impairment of gene regulation^1^. These mutations may ultimately jeopardize the integrity of the host genome. Therefore, the present-day host-TE interactions are the only ones that did not lead to extinction of either the host or the TE^2^ Unraveling the diversity of such host-TE interactions is a pivotal area of research that provides valuable insights into genome function and disease biology.

TEs generally remain within the colonized host genome, except for rare instances of horizontal transfer^3^. To ensure its survival, each TE must anticipate its progressive mutational decay by inserting new functional copies into the host germline DNA, allowing vertical transmission to next generations. However, the rates of such replicative transpositions have been set, during evolution, at low levels compatible with the survival of both the host and TEs. Some of the mechanisms controlling TE transposition involve a specific class of small regulatory RNAs known as Piwi-interacting RNAs (piRNAs), which, when associated with PIWI proteins, a subclass of Argonaute proteins, can hybridize with nascent or cytoplasmic TE transcripts. This specific targeting by the host defense machinery leads to the silencing of TEs, either transcriptionally (TGS) or post-transcriptionally (PTGS) respectively^4^.

In *Drosophila melanogaster*, comparative analyses of the conserved reverse transcriptase domain of Long Terminal Repeat RetroTransposable Elements (LTR-RTEs), which replicate via an RNA intermediate, revealed that they are distributed into three clades: Copia, BEL and Gypsy^5,6^. LTR-RTEs of the Copia and BEL clades each encode a single open reading frame (ORF) and are represented by species a few species as compared with the many species of the Gypsy clade species also displaying a stronger heterogeneity in coding sequence, with one, two or three ORFs^5,6^. Several TE species of the latter clade are specifically expressed in the somatic cells surrounding the germline, their replication being dependent on the ability to infect germline cells. This infectious capacity, that is linked to the acquisition of an ORF encoding a viral-like envelope in their common ancestor^5,7^, is assumed to have preserved the host germline by starting the replication cycle in the adjacent somatic tissue and therefore to have been responsible for the evolutionary success of this clade. Indeed, each ovarian somatic cell type has been exploited as a TE-specific expression niche^7^, free of competitive interactions and interference between copies of related TE species. This niche partitioning probably arose after more or less successful attempts by the TE to co-opt endogenous co-factors for transcriptional regulation.

At the integration step of the TE replication cycle, co-optation of endogenous proteins as co-factors to specifically insert into either neutral or less essential regions of the genome, is thought to result in tolerant integration niches. Indeed, in organisms with small gene-rich genomes like *S. cerevisiae* and dictyostelids, the insertion into, or close to, coding genes being mutagenic, some TEs preferentially target regions considered as less critical, such as gene-poor heterochromatin or redundant, non-essential multicopy genes like tDNA and rDNA^8^. However, since sharing the same neutral insertion site may lead to competition between TEs, each TE is expected to exhibit its own insertion site preferences. Indeed, recent data suggest that P-elements favor replication origins in Drosophila while LTR-RTEs preferentially integrate near promoters and exons of active genes^9^. Whether integration preferences vary between different LTR-RTE species and/or are influenced by specific cellular contexts requires further investigation.

Studying TE ecology regarding not only the interactions between a TE and its host but also between members of the whole community of TE species having colonized the host (the set of TE copies coexisting in the genome of the host) is expected to provide further insights into the ways TE have successfully invaded all present-day eukaryotic genomes.

To investigate LTR-RTE ecology, we used a previously constructed *Drosophila melanogaster* line that allows inducible impairment of LTR-RTE repression and then determined the timing and specificities of LTR-RTE expression and integration. Using short-read genome resequencing, we had previously observed a few putative insertions for two LTR-RTE species of the Gypsy clade (gtwin and ZAM) following this de-repression, providing a proof of concept for this approach^10^. At the time, the localization of newly integrated LTR-RTEs was not possible, and the number of integrations was not sufficient to perform a comprehensive comparison of the insertional landscapes.

Here, using long-read sequencing to annotate the genomes of flies that had been subjected to LTR-RTE de-repression for many successive generations, we observed the accumulation of LTR-RTEs from different clades. At the 73rd generation, we mapped enough newly inserted copies of four LTR-RTEs (a total of 798 insertions of roo, copia, gtwin or ZAM) to disclose specific biases in their landing site preferences. Indeed, since natural selection had only modestly altered the landscape of these recently integrated LTR-RTEs, it had not yet erased the memory of their initial preferences for insertion sites. Moreover, gtwin and ZAM exhibited differences in expression patterns, but also in the timing of their integration into the embryonic genome, leading to different landing site preferences. Our findings highlight how, over the course of evolution, the diverse cell identities exploited by various LTR-RTE species for both expression and integration have led to the colonization of TE-specific genomic niches of integration.

## Results

### Four LTR-RTEs integrate into the Drosophila germline after successive generations of somatic Piwi knockdown

To accumulate a substantial number of LTR-RTEs in the *Drosophila* genome, we used a strain previously constructed in our laboratory^10^. In this strain, a traffic-jam-Gal4/Gal80 inducible driver activates, at high temperature, the expression of a short RNA hairpin construct targeting Piwi (sh-piwi) in the gonadal somatic cells. This somatic knockdown (sKD) alleviates LTR-RTE repression in these cells without causing sterility^10^. When females containing traffic-jam-Gal4/Gal80>sh-piwi are shifted, for a few days, from the 20°C non-permissive to the 25°C permissive temperature, they display a partial depletion of the Piwi protein in their ovarian somatic cells (piwi-sKD), leading to an accumulation of LTR-RTE transcripts in the ovaries and *de novo* transposition^10^. As previously shown, mobilization of LTR-RTEs already occurs after the induction of Piwi depletion in a single generation^10^. Here, we first confirmed that the fertility of shifted flies and of their progeny were not sufficiently affected to prevent the induction of piwi-sKD during successive generations (Figure S1).

At the 11^th^ (G11), 31^st^ (G31) and 73^rd^ (G73) generations, we isolated a large subset of the shifted population to maintain a stock constantly at 20°C, the non-permissive temperature for piwi-sKD (Figure 1A). Additionally, as a negative control, we maintained a fraction of the initial parental line G0 constantly at 20°C for 100 generations (G0F100). This experimental scheme provided us with the G0F100 control population that had not undergone any temperature shift, along with the G11, G31 and G73 populations with increasing occurrences of piwi-sKD and hopefully harboring an accumulation of new LTR-RTE insertions in their respective genomes.

**Figure 1:**
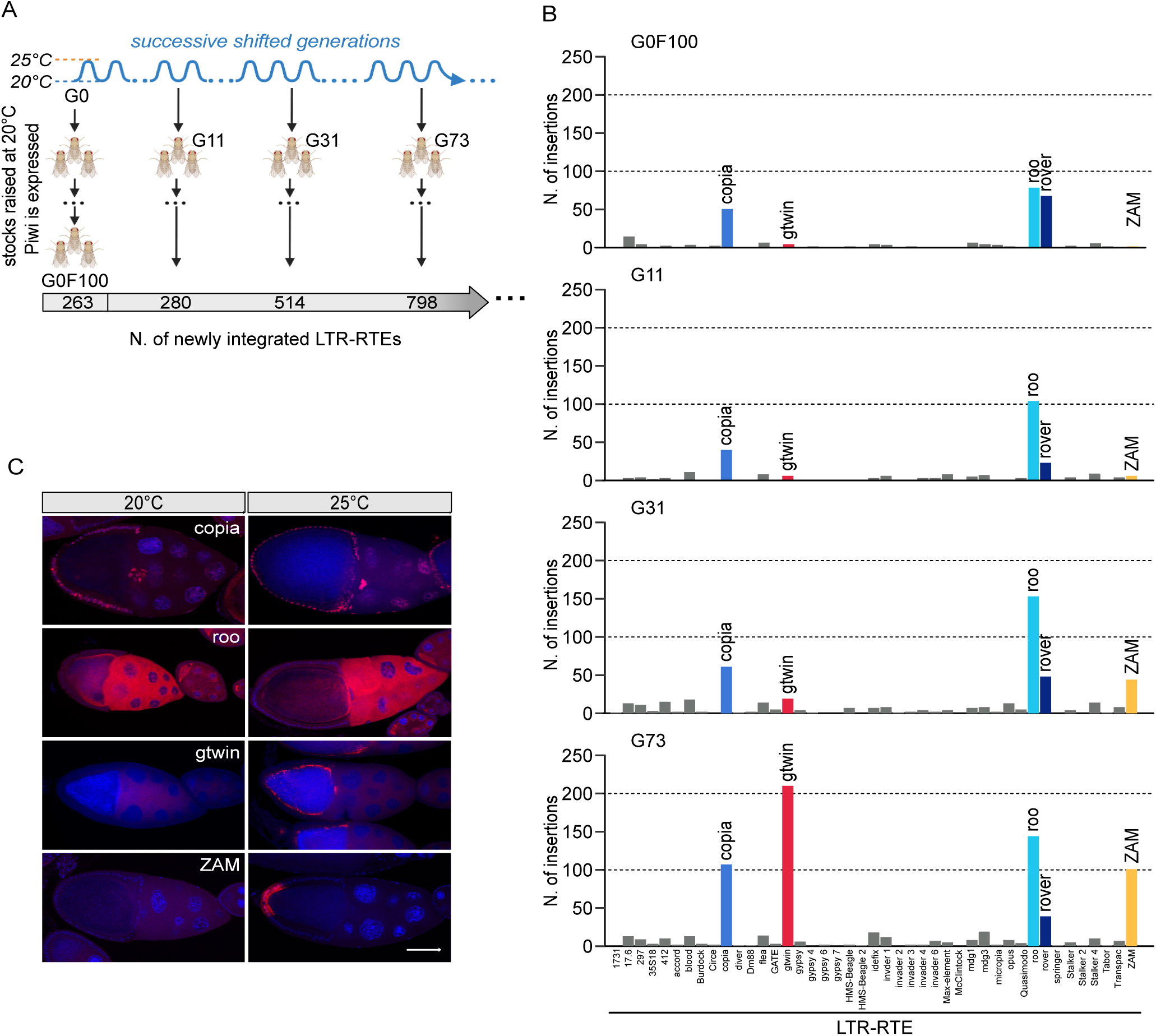
Four LTR-RTEs integrate into the Drosophila germline after successive generations of somatic Piwi knockdown. (A) Schematic representation of the successive transient Piwi somatic knockdowns (Piwi-sKD), induced by shifting adults from 20°C to 25°C for 5 days in each generation, followed by constant maintenance at 20°C of three large populations isolated from generations 11, 31 and 73. G0F100 corresponds to the initial G0 parental line that was constantly kept at 20°C for 100 generations. Bottom: The grey arrow contains the numbers of new LTR-RTE insertions detected in the different isolated populations as compared to the initial G0 parental line. (B) Quantification of the new LTR-RTE insertions annotated in the control generation (G0F100) and the successive shifted generations G11, G31 and G73, as compared to the initial parental G0 line. (C) Representative images of stage 10 ovarian expression patterns obtained for copia, roo, gtwin and ZAM LTR-RTEs by smiFISH (in red) at the 20°C restrictive temperature and after a 5-day-adult shift to the 25°C permissive temperature. DNA was stained with with 4,6-diamidino-2-phenylindole (DAPI; blue). Bar represents 50 µm.

Using the Oxford Nanopore Technology (ONT) for long-read sequencing of DNA, we sequenced the genomes of 100 pooled males originating from the G0, G0F100, G11, G31, and G73 populations (Figure 1A). We used the TrEMOLO pipeline^11^ to annotate the newly integrated LTR-RTEs. This bioinformatics tool aligns and compares long-read sequences obtained at each defined generation with those of the initial G0 parental genome (Materials and Methods). We identified an increasing number of new LTR-RTE insertions in the successive shifted generations: with 280, 514, and 798 new insertions for the G11, G31, and G73 generations respectively (Figure 1A). As shown in Table S1, the majority of these newly identified LTR-RTE insertions, which belong to 43 species, rarely reached 20 new insertions per species. This could result from a relatively low expression of some of these LTR-RTEs together with a low transposition rate and/or a high selective pressure against new insertions. Nevertheless, among the different LTR-RTE species, five (namely roo, copia, rover, ZAM and gtwin), covering the three known clades of LTR-RTEs (Copia; Gypsy; BEL), had equal or more than 40 new insertions in the successive shifted populations (Figure 1B, Table S1).

Through the same bioinformatics analysis of the G0F100 long-read sequences, we detected 263 insertions that were absent from the G0 initial parental genome (Figure 1A, B). They uniquely corresponded to roo, copia and rover, suggesting that these three LTR-RTEs were able to express and transpose spontaneously (independently of piwi-sKD treatment). To confirm that these three elements were indeed expressed in the G0F100 stock, we assayed their expression patterns using single molecule inexpensive fluorescent *in situ* RNA hybridization (smiRNA-FISH) at 20°C and 25°C (Figure 1C). These observations confirmed that roo and copia are indeed expressed regardless of temperature and revealed specific expression patterns for these two elements in the ovaries of G0F100 females. More specifically, roo was expressed in the germline and its transcripts accumulated in the oocyte cytoplasm, similar to a LINE element like the I-element^12,13^. We detected transcripts of copia in the nuclei of the follicle cells that surround the germline. The same analysis for rover expression was inconclusive, as we did not detect transcription in shifted or non-shifted ovaries (data not shown). We hypothesized that most of the new rover insertions in the different populations resulted from somatic transposition occurring during development. In agreement with this possibility, we quantified the frequency of the different new rover insertions and found it was always low (Table S2). Moreover, we could not obtain any evidence for vertical transmission of these rover insertions. Indeed, in contrast to some copia and roo insertions shared between the successive populations and therefore likely to correspond to germline inherited insertions, we could not detect a single *rover* insertion that was shared at the same genomic position by at least two generations (Table S3).

Contrary to the aforementioned LTR-RTEs, ZAM and gtwin, belonging to the Gypsy clade, had exclusively transposed in the shifted generations, as demonstrated by their increased numbers of insertions (Figure 1B). As this suggests that their expression was regulated by the presence of Piwi protein in the follicle cells, we analyzed their expression by smiRNA-FISH at restrictive and permissive temperatures. These two elements are clearly not expressed in the ovaries at 20°C and start to express in follicle cells of the ovaries at 25°C, when Piwi has been depleted (Figure 1C). Interestingly, these two elements exhibit specific niches of expression that are selective to the somatic follicle cells. ZAM expression was restricted to posterior follicle cells as previously reported^7,14^, whereas gtwin seemed to have a broader expression pattern throughout the follicle epithelium (Figure 1C).

With the deep analysis of long reads obtained by ONT sequencing and the assembly of the G0 genome, we could identify three full-length copies of gtwin, two being localized on chromosome Y and one on chromosome 2R. For ZAM, we identified at least two full-length ZAM copies located on chromosome 2R. Each of these species of LTR-RTEs had produced 6 new insertions after 11 generations of Piwi depletion, and 44 (ZAM) and 19 (gtwin) after 31 generations. This trend intensified with successive generations, with 101 and 210 new insertions in the 73^rd^ generation, respectively (Table S1).

Altogether, our long-read sequencing analysis of several generations, submitted to successive somatic Piwi depletion in ovaries, revealed that four LTR-RTE species were able to move in the *D. melanogaster* genome and thereby generate a sufficient number of germinal insertions to warrant further studies.

### Selection had little impact on the landscape of newly integrated insertions

Using the new insertions annotated in G73, we investigated whether specific insertion sites were favored by each of the four active LTR-RTE species. Aware of the potential biases introduced by selection on the TE insertion landscapes^15^, we first tried to assess the extent of such additional biases in the G73 dataset. To do so, we segmented the *D. melanogaster* genome into intergenic regions, introns, and exons, assigned the new insertions found in G73 to either of these three bins, and statistically compared the observed-to-expected numbers under the null hypothesis of random insertion (Table S4). For each of the four species of LTR-RTEs (copia, roo, gtwin and ZAM), the observed numbers of intergenic insertions were roughly in line with expectation (Figure 2A). This suggests that the previously reported purifying selection operating over longer periods against genic insertions^9,16^ did not have enough generations to introduce such a bias in our sequencing data. Moreover, we detected only 2 times less insertions in exonic regions than expected in G73, whereas a tenfold reduction compared to expected was observed for much older insertions^9^. Interestingly, we also detected more insertions in intergenic regions, whereas 1.2-fold depletion was noted for older insertions^9^.

**Figure 2:**
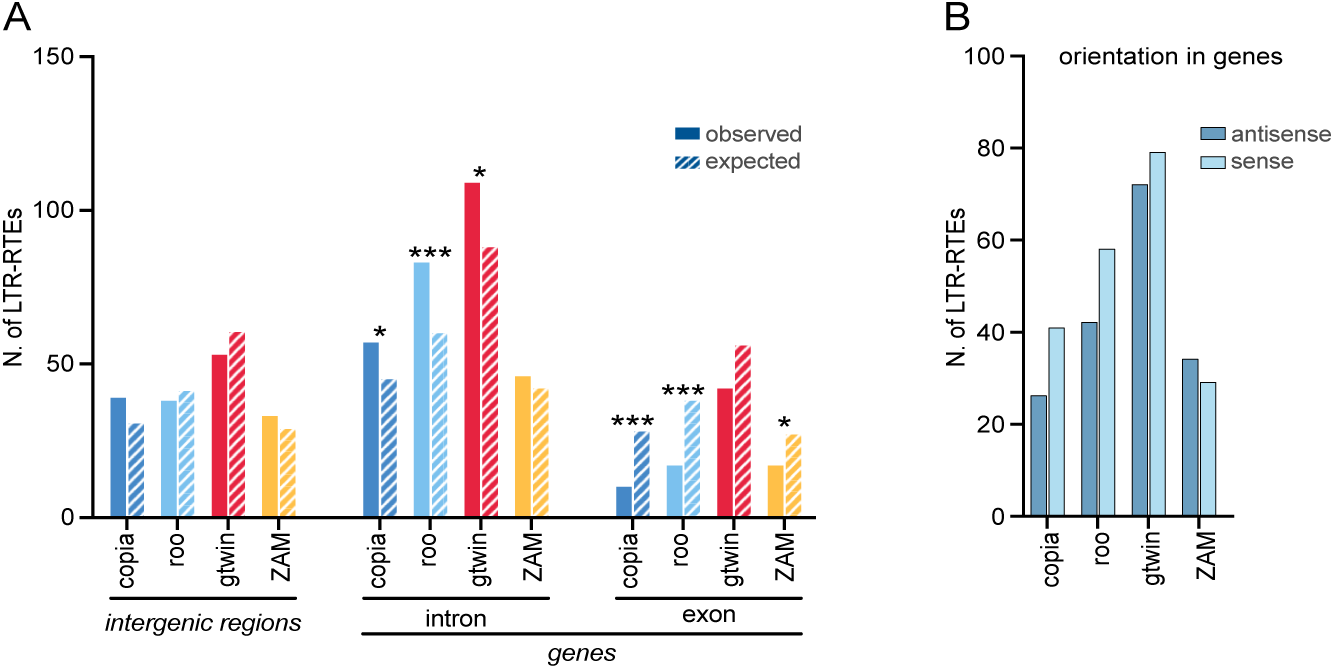
Selection had little impact on the landscape of newly integrated insertions. (A) Barplots depicting the observed (filled) and expected (dashed) numbers of LTR-RTEs in the intergenic, intronic and exonic regions of the genome. Expected values were calculated according to the size of each genomic region considered and the total number of new insertions obtained for each LTR-RTE. p-values were calculated using binomial tests corrected by Benjamini–Hochberg step up procedure to control the false discovery rate; p-value: * <0.05, ** <0.005, *** <0.001. (B) Numbers of insertions for each LTR-RTE that had occurred into the antisense strand (dark blue) or sense strand (light blue) of protein-coding genes. No significative differences were observed between insertions in these two orientations using binomial tests corrected by Benjamini–Hochberg.

A second feature of selection on the TE insertion landscape is the elimination of genic insertions that are oriented in the sense direction of transcription. It is believed that insertions oriented in the sense direction are more detrimental than those in the antisense direction, as the latter are less likely to influence gene transcription signals^17^. In G73, we did not notice any significant difference in the number of insertions in these two opposite orientations and rather a higher number of insertions of the different LTR-RTEs in the sense orientation. Selection over 73 generations had apparently not lasted long enough to create the expected orientation bias in our population (Figure 2B). Finally, in the G73 population, we did not detect any signs of positive selection for newly integrated LTR-RTEs, as the vast majority were present at very low frequencies within the population (Figure S2A). More particularly, none of the LTR-RTEs newly integrated in piRNA clusters, showed any tendency to invade the G73 population (Figure S2B and S2C). Altogether, we were confident that selection did not have enough time to significantly affect the landscapes of our newly inserted LTR-RTEs.

### Each LTR-RTE species has its own specific chromatin states preferences for genomic integration

Eukaryotic genomes are partitioned into chromatin domains containing different epigenetic states that are essential for proper gene regulation^18,19^. So far, insertions of various TEs have been mainly associated with open chromatin structure^20,21^, suggesting that a higher accessibility of genomic DNA could help TE integration. To test whether a higher degree of chromatin flexibility is a driver of integration site-specificity, we analyzed whether LTR-RTE insertions were preferentially associated with some of these different chromatin states. We compared the distributions of the LTR-RTE insertions observed in the G73 population with those expected from the relative genomic proportions of the nine distinct chromatin states defined in S2 cells^18^. These cells are derived from late male embryonic tissues (stages 16-17). While we detected insertions of the different species within all nine chromatin states, we mainly observed significant enrichment in chromatin states 1 to 4 characterized by their openness relative to the other chromatin states (Table S4) (Figure 3). Each of the four active LTR-RTE species had its own specific pattern of insertions. This specificity was particularly evident for gtwin insertions that showed a significant enrichment in chromatin states 1 and 3, corresponding to promoter- and enhancer-like chromatin, respectively (Figure 3). In contrast, copia insertions showed a broader distribution, with significant enrichment in chromatin states 3, and higher-than-expected detection in states 4 and 7. roo and ZAM rather accumulated in the 4th chromatin state defined as open chromatin with enhancer features including H3K36me1 but devoid of H3K27ac (Figure 3)^22^. Overall, the distinct profiles obtained for LTR-RTE insertions in G73 are not the result of passive processes but could be driven by active integration mechanisms that contribute to target different LTR-RTEs to specific states of open chromatin.

**Figure 3:**
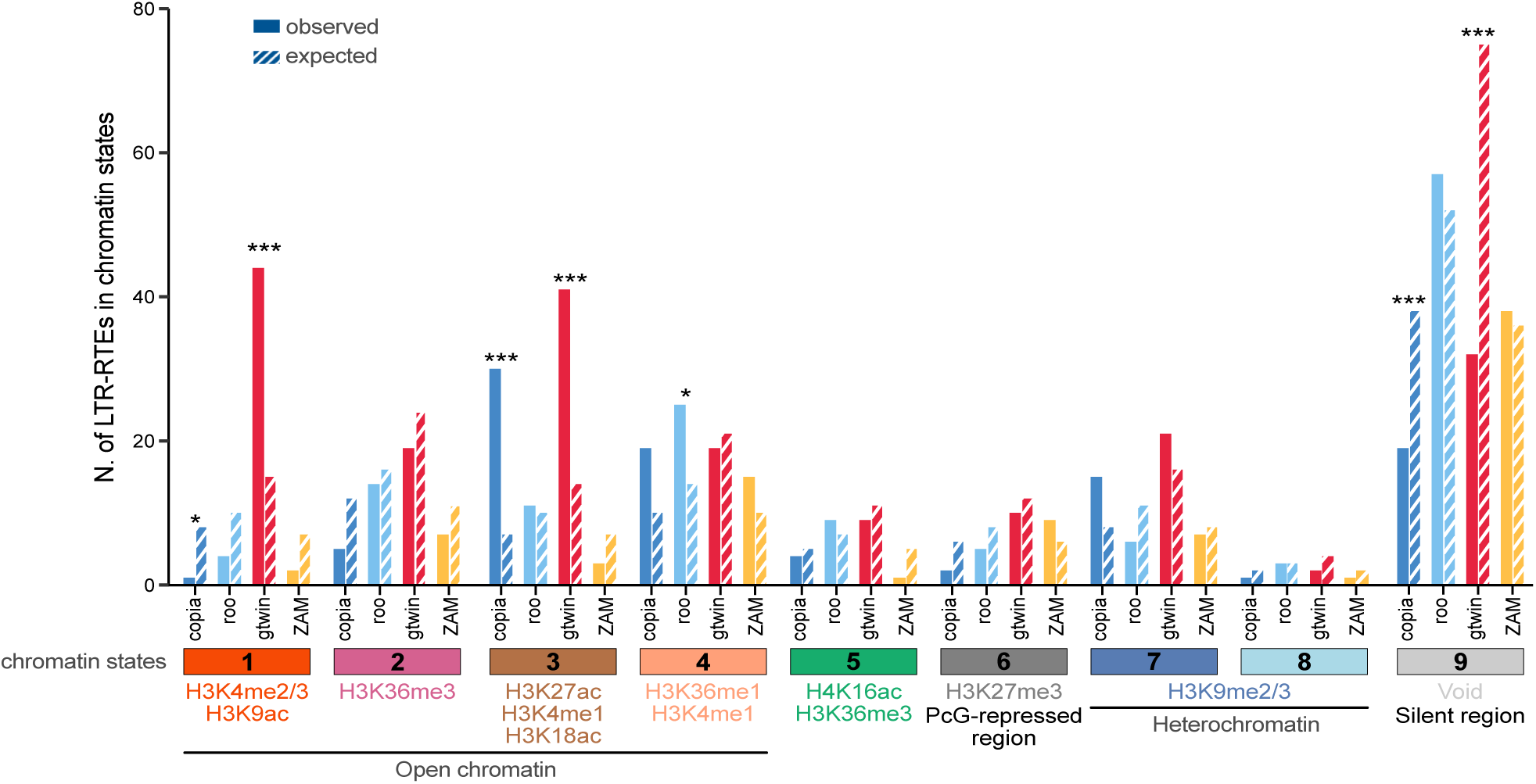
Each LTR-RTE species has its own specific chromatin states preferences for genomic integration. Barplots depicting the observed (filled) and expected (dashed) numbers of LTR-RTEs in the nine described chromatin states. Typically enriched post-translational histone marks in each chromatin state are indicated below the corresponding state. Expected values were calculated according to the size of each chromatin state in the genome and the total number of new insertions obtained for the indicated LTR-RTEs. Statistical significances (p-values) were calculated using binomial tests corrected by Benjamini–Hochberg step up procedure to control the false discovery rate; p-value: * < 0.05, ** < 0.005, *** < 0.001.

### gtwin preferentially inserts into the open chromatin of enhancer and promoter regions before cellularization

We then focused on the timing of the germline integration of gtwin and ZAM, the only two enveloped LTR-RTEs derepressed in the ovarian follicle cells: how long does this integration step occur after the expression step? Indeed, previous genetics studies of gypsy/mdg4, a similar infectious LTR-RTE with an envelope-coding gene, have indicated that, despite expression during oogenesis, integration occurs only in the genome of the progeny^23^. Such a long delay of the integration step might correlate with the egg laying-induced decondensation of the oocyte karyosome. We therefore expected that ZAM and gtwin would also integrate during embryogenesis.

Given our findings that gtwin insertions were significantly enriched in promoter- and enhancer-like open chromatin states 1 and 3^18^, we focused on the corresponding embryonic epigenetic marks. We leveraged available modENCODE Chromatin ImmunoPrecipitation data of the relevant histone marks coupled with high-throughput sequencing (ChIP-seq) at late (16 to 20 hours after egg laying (AEL))^24^ and early (∼2.5 hours AEL, corresponding to stage 5)^25^ embryogenesis. In this analysis, ZAM was used as a negative control as it is rather associated with chromatin state 4, an open chromatin state that lacks the four tested histone modifications^18,22^. This analysis suggested that gtwin insertion sites tend to be enriched in H3K4me3/H3K9ac and/or H3K4me1/H3K27ac epigenetic histone marks already deposited in 2.5-h embryos and likely corresponding to promoter and enhancer regions, respectively (Figure 4A).

**Figure 4:**
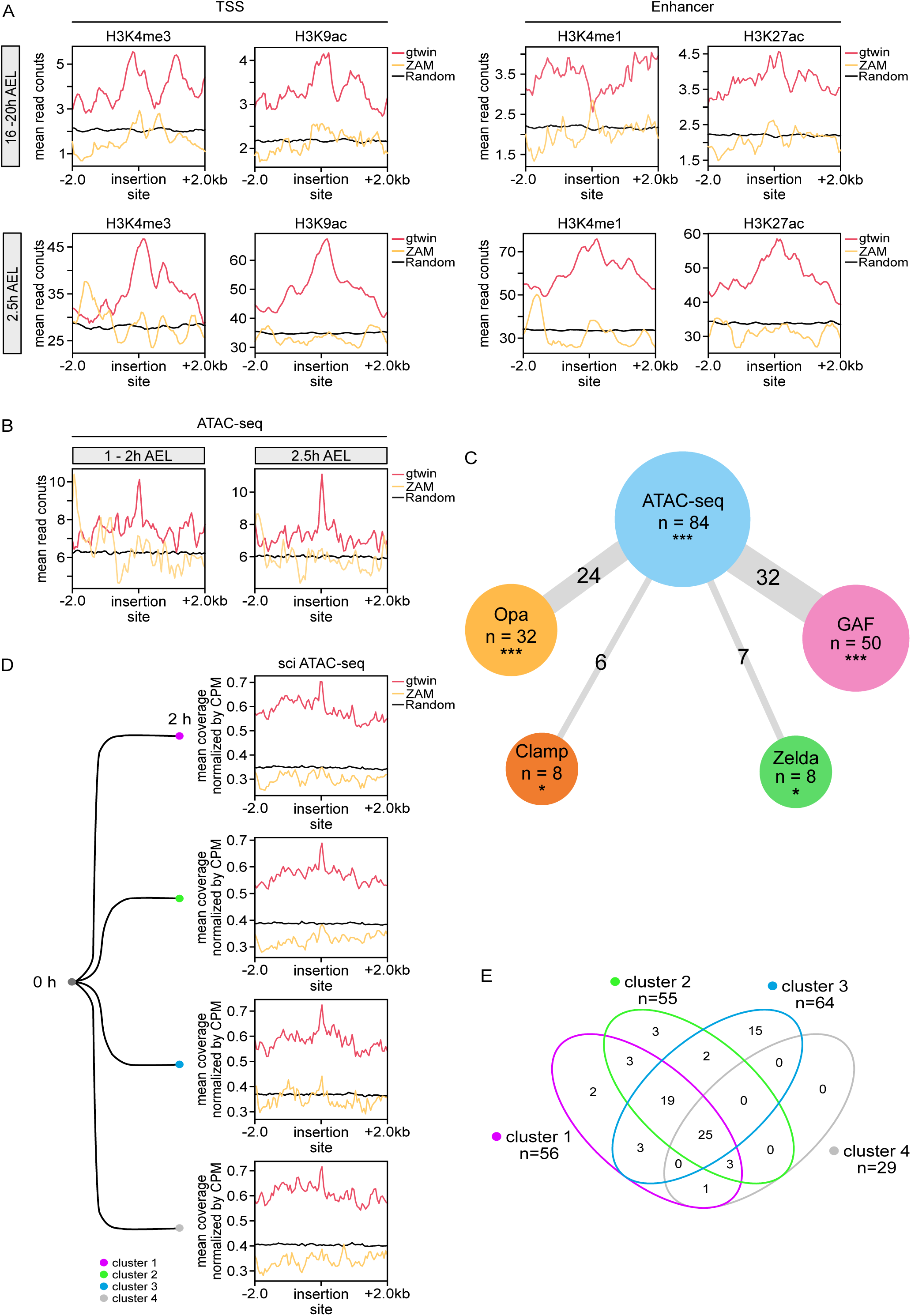
gtwin preferentially inserts into the open chromatin of enhancer and promoter regions before cellularization. (A) Metaplots depicting the mean read counts of ChIP-seq data for H3K4me3, H3K9ac, H3K4me1 and H3K27ac obtained in embryos 16-20 h (top panels) and 2.5 h (bottom panels) after egg laying (AEL). 4-kb windows centered either on the 210 gtwin (red curve) or on the 101 ZAM (yellow curve) novel insertions found in G73 population. Black curve (random) was obtained by averaging the mean read count values of 100 positions randomly selected 100 times on the same genome. (B) Metaplots depicting the mean read counts obtained by ATAC-seq on embryos 1-2 h (left) and 2.5 h (right) AEL, considering the same 4 kb windows as in A. (C) Overlap of pioneer-factor-rich and ATAC-seq-rich gtwin insertion sites in stage 5 embryos. The size of each circle is scaled by the number of gtwin insertions that are enriched for accessible chromatin (blue), GAF (pink), Zelda (green), Clamp (orange) and Opa (yellow). The number indicated in the edges corresponds to the number of gtwin insertions possessing both features. Expected values were calculated according to the proportion of ChIP-seq signal of the protein of interest and the total number of new gtwin insertions. P-values were calculated using binomial tests corrected by Benjamini–Hochberg step up procedure to control the false discovery rate; p-value: * < 0.05, ** < 0.005, *** < 0.001. (D) Metaplots depicting in 4 kb window centered on 210 gtwin (red), 101 ZAM (yellow) or 100 random insertion sites (black), single-cell ATAC-seq (sci ATAC-seq) mean normalized coverage (by Counts Per Million unique mapped reads) of the 4 nuclei clusters identified in 0-2 h embryos. (E) Venn diagram highlighting the distribution of the 76 gtwin insertions sites enriched in sci-ATAC-seq signal among the 4 clusters.

Interestingly, these regions have been described as among the first to become open and accessible from stage 3-4 (1 to 2 hours AEL)^26^. Focusing on chromatin accessibility, by using previously published ATAC-seq data from whole early embryos^27^, we confirmed that gtwin insertion sites were preferentially localized in the open chromatin peaks of stage 3-4 and stage 5 embryos (1-2 and ∼2.5 hours AEL, respectively) (Figure 4B). By contrast, ZAM insertion sites did not correspond to chromatin regions that are open at these early stages of embryogenesis. This open chromatin state, with low nucleosome occupancy and few higher-order chromatin structures^28^, is believed to result from the binding of a special class of transcription factors known as pioneer factors. These pioneer factors are unique in their ability to overcome nucleosomal barriers that establish accessibility of cis-regulatory elements required for further DNA transactions^29,30^. We wondered whether the binding of a specific pioneer factor could be correlated with the gtwin insertion sites. To investigate this, we identified the gtwin insertions located in open chromatin of stage 5 embryos (n=84) and then analyzed the stage 5 embryonic ChIP-seq data of four pioneer factors: GAGA Factor (GAF), Opa, Chromatin-linked Adaptor for MSL proteins (CLAMP), and Zelda ^31–34^. As expected, most of the stage 5 open chromatin regions containing a gtwin insertion (69 out of 84) were bound by at least one of these pioneer factors. However, it appears that no specific pioneer factor was responsible for gtwin binding: only 32, 24, 6 and 7 of these regions correlated with GAF, Opa, CLAMP and Zelda binding sites, respectively (Figure 4C). Altogether, our analyses revealed that chromatin accessibility, independently of the pioneer factor associated, is the most significant feature determining the choice of gtwin insertion sites in early embryos.

Using this preferential chromatin feature associated with gtwin integration, we wondered whether it would be possible to determine its timing of integration during embryogenesis. Notably, we wanted to determine if germline gtwin integrations had occurred before or after germline specification, a process that naturally initiates around 1.30h AEL^35^. To do so, we took advantage of single cell indexing sci-ATAC-seq data which, during successive two-hour intervals of Drosophila embryogenesis, enables clustering of individual nuclei on the basis of similarity of their chromatin accessibility patterns^36^. We determined the averaged chromatin accessibility within 4 kb windows around gtwin insertion sites in the four first ATAC-seq clusters of nuclei identified at 0-2 hours AEL. For each cluster, gtwin insertion sites tended to be localized in open chromatin (Figure 4D). From peak calling we estimated that 76 gtwin insertions were present in regions open in at least one of the four clusters studied at 0-2 hours AEL. Interestingly, 56 of these insertion sites were present in at least two clusters (Figure 4E). This suggests that most gtwin insertions occurred during very early syncytial stages of Drosophila embryogenesis, before the differentiation of these first clusters and probably also before the specification of germ cells, the nuclei of which being the first to become cellularized at this stage^35^.

### Chromatin accessibility of ZAM insertion sites correlates with ZAM late embryonic expression

Our analyses of chromatin accessibility did not reveal any preference for ZAM integration in open chromatin regions during early embryonic development (Figures 4B, D). To investigate the chromatin accessibility of ZAM insertion sites in later developmental stages, we used time-course sci-ATAC-seq data available at 2-hour intervals spanning the entire embryogenesis^36^. Across eight intervals from 0 to 16 h AEL, we calculated the average sci-ATAC-seq signal for 101 ZAM insertions and 210 gtwin insertions, normalized with signals from 100 randomly selected insertions (Materials and Methods) (Figure 5A). This analysis confirmed that gtwin insertion sites already exhibited an accessible chromatin status in the very early stages of embryogenesis that was then maintained at that high level throughout embryonic development. Concerning the status of the ZAM insertion sites, the chromatin gradually became more accessible during embryogenesis, particularly in 10-12 h, 12-14 h, and 14-16 h AEL embryos (Figure 5A). Altogether this temporal analysis suggested that ZAM might integrate later during development.

**Figure 5:**
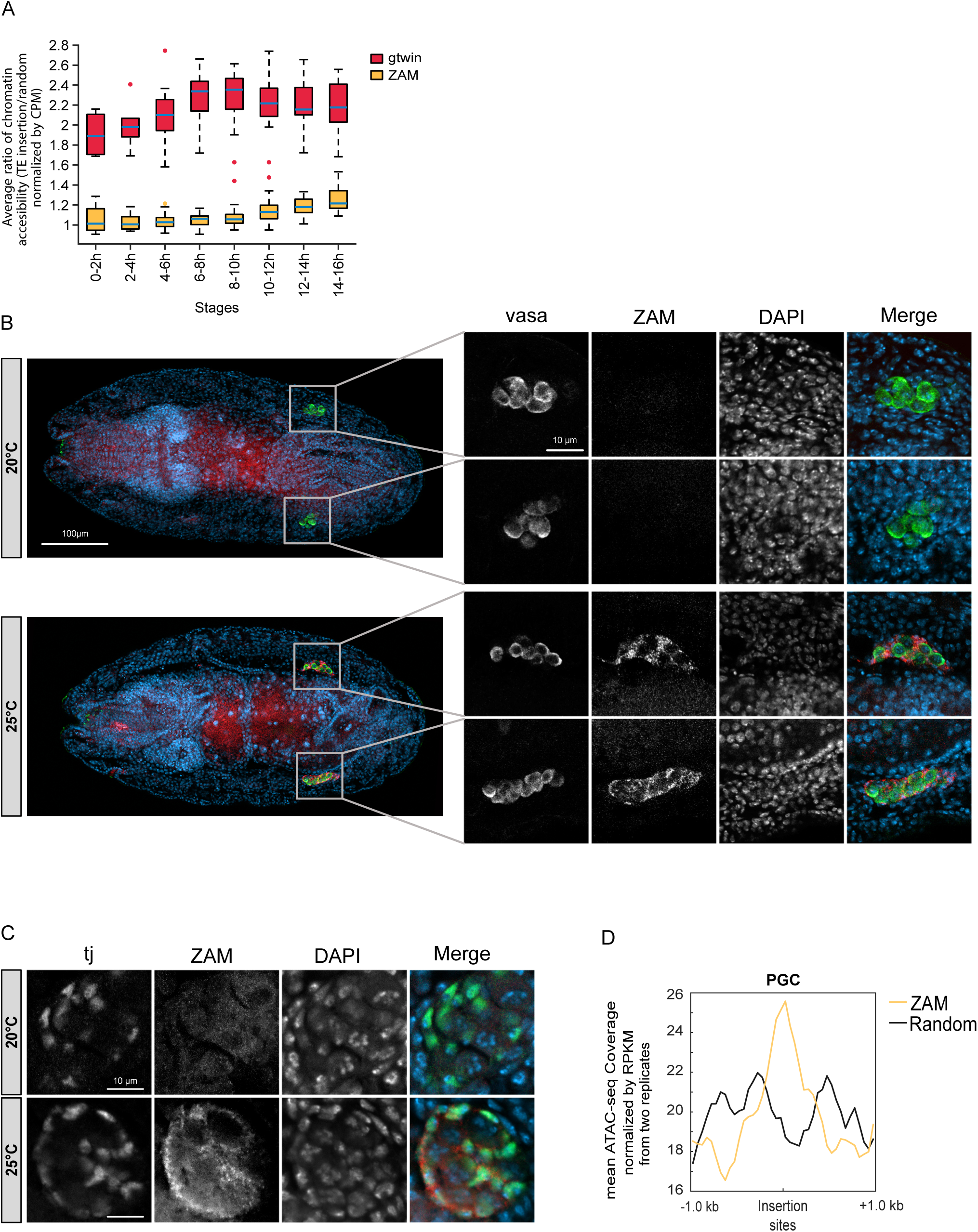
Chromatin accessibility of ZAM insertion sites correlates with ZAM late embryonic expression. (A) Boxplots representing the temporal kinetics of the average chromatin accessibility around gtwin (red) and ZAM (orange) insertions, relative to random profiles. At each time window is presented the distribution of the relative chromatin accessibility computed for the set of defined cell clusters. At each time window and for each cell cluster, this accessibility is defined as the average sci-ATAC-seq signal of the 200 bp windows centered on LTR-RTE insertions divided by the average signal of 100 randomly selected 200 bp windows. (B) Vasa immunostaining combined with ZAM smiFISH on 12- to 16-h whole-mount embryos at 20°C (upper panel) and 25°C (lower panel). The right panel shows higher magnification of embryonic gonads. Anti-vasa antibody (green) labelled the primordial germ cells (PGCs) of the gonads. ZAM transcripts labelled in red are detected in cells surrounding vasa-positive cells at 25°C (lower panels). DNA is labelled with 4,6-diamidino-2-phenylindole (DAPI; blue). (C) Traffic jam immunostaining combined with ZAM smiFISH on gonadal cells of 12- to 16-h embryos at 20°C (upper panel), 25°C (lower panel). The somatic primordial gonadal cells (SPGs) labelled in green are tj-positive cells. ZAM transcripts labelled in red are detected in tj-positive cells at 25°C (lower panel). DNA is labelled with 4,6-diamidino-2-phenylindole (DAPI; blue). (D) Metaplot showing, in 2kb windows centered on 101 ZAM (yellow) or 100 random insertion sites (black), the mean ATAC-seq signals of two replicates, normalized by coverage (by Counts Per Million unique mapped reads: RPKM), of PGCs purified from overnight embryos.

Two non-exclusive hypotheses may explain the delayed timing of ZAM integration into the germline. The first possibility is that ZAM viral particles infecting the germline during oogenesis remain dormant during the early stages of embryogenesis. Alternatively, ZAM might undergo a second wave of expression at later stages of embryogenesis. Since the eggs were collected at 25°C, a temperature that permits piwi-sKD, a second wave of ZAM de-repression could occur. To test this possibility, we combined ZAM smiFISH and germline-specific vasa immunostaining in late embryos. As shown in Figure 5B, we observed a high expression of ZAM in the gonads of the embryos laid at 25°C but not at 20°C. At these late stages of embryogenesis, the primordial germ cells (PGCs) have migrated away from the midgut toward the adjacent mesoderm and have become associated with somatic gonadal precursors (SGP)^35,37^ that express traffic jam (tj)^38^. As expected, we observed ZAM expression in the tj*-*positive cells of the late embryonic gonads specifically at 25°C (Figure 5C). Moreover, the expression of ZAM in SGPs at 25°C is consistent with the permissive temperature for piwi-sKD, as piwi-sKD is driven by traffic-jam-Gal4 at 25°C and not at 20°C due to the presence of Gal80ts. Overall, these analyses show that ZAM is expressed at later stages of embryonic development, in the SGP cells positive for tj and sensitive to Piwi depletion when embryos are laid at 25°C.

To determine whether this later timing of ZAM expression in embryos could result in germline-specific insertions, we decided to determine whether ZAM insertion sites correlate with chromatin regions that are accessible in the late embryonic germ cells. To do so, we took advantage of a *Drosophila* line expressing a GFP tagged version of the germline-specific gene *vasa*^39^ to perform GFP cell sorting using FACS^40^ (Materials and Methods) from cell extracts originating from overnight egg collections. After cell sorting, ATAC-seq experiments were performed in duplicates on embryonic GFP-positive cells corresponding to PGCs. We compared the averaged profiles of the ATAC-seq signals, performed in duplicate, in the 2kb windows centered around the 101 ZAM insertions sites, with those obtained for 100 randomly chosen 2kb windows in the same cells. We observed that in the PGCs, the averaged ATAC-seq profile reached a maximum centered on the ZAM insertion sites (Figure 5D). Altogether these experiments show that ZAM insertion sites correlate with a specific chromatin accessibility landscape in late PGCs that was not observed in the four early embryonic clusters before cell differentiation (Figure 4D). Overall, we propose that the late wave of ZAM somatic expression could lead to PGC infection, and the subsequent targeting of ZAM integration machinery to genomic regions that are accessible in late PGCs.

## Discussion

Transposable elements may alternate rapid bursts of activity and prolonged phases of repression during which these genomic parasites do not replicate efficiently within the host genome^41^. Present-day TE-TE and TE-host genome interactions are probably the outcome of multiple co-evolutions, allowing the peaceful coexistence of different TE species within the same genome. In an attempt to describe the diversity of these interactions, we took advantage of a particular *Drosophila melanogaster* laboratory strain^10^ to simultaneously impair the repression of, and therefore awaken, several LTR-RTEs. We observed that four active elements (roo, copia, gtwin and ZAM) had been able to efficiently transpose into the Drosophila germline, after several dozens of successive generations of LTR-RTE de-repression. We found that they all differ in several steps of their replication cycles.

Regarding the expression step, even though they were all transcribed at the end of oogenesis, the RNAs of one of them, roo, were specifically expressed in the germinal nurse cells, while those of the three others were found in different cell-types/compartments of the somatic follicular epithelium, either at the posterior pole, for ZAM, or ubiquitously for gtwin and copia (but, in the latter case, sequestered into the nuclei). We also disclosed a second window of ZAM somatic expression in the SGPs of the late embryonic gonad.

Concerning the integration step, our approach was based on characterizing the overall epigenetic specificity of their genomic insertion sites. We thus found that, although each of the four landing site landscapes correlated with open chromatin, they all seemed to display specific epigenetic preferences. Copia and roo insertions were found predominantly in chromatin states 3 and 4, respectively, while gtwin appeared to preferentially choose promoter- and enhancer-like landing domains, enriched in chromatin states 1 and 3, and ZAM would rather favor chromatin state 4. Moreover, our data indicated that the preference for specific genomic insertion sites may follow the differentiation of the chromatin landscape of the cells that are invaded by gtwin and ZAM at different stages of the embryonic development. Indeed, for maternally deposited gtwin, a significant proportion of the insertions seemed to have occurred as soon as their landing sites had begun to be accessible, at the very beginning of embryogenesis. Conversely, consistent with the late embryonic wave of ZAM somatic expression, several ZAM insertions were located within different open chromatin regions, accessible only in late embryonic germ cells. However, although the insertion of a LTR-RTE into closed chromatin is generally considered unlikely, it cannot be entirely ruled out. Therefore, we cannot exclude the possibility that some maternally deposited ZAM virus-like particles would have driven integration at early stages into close chromatin landing sites that would open later in the gonadic PGCs. Altogether, our findings disclose a novel level of LTR-RTE niche partitioning linking temporal and spatial features of the integration step of the replication cycle.

### Diversity of expression and integration niches

The diversity of ovarian expression patterns reported here (Figure 1) has also been observed recently for 16 species of evolutionarily related LTR-RTEs^7^. These different patterns in the onset of the replication cycles of this TE class, indicate that each LTR-RTE species has evolved specific interactions hijacking tissue-specific transcription factors to adapt their proper expression niche to a specific cell type. In our study, we identified a novel cell type in which ZAM is also expressed, the SGPs of late embryonic gonads. Future experiments will be necessary to determine whether the transcription factor called Pointed, which drives ZAM transcription in the posterior part of the ovarian follicular cells^42,43^, is also responsible for its expression in the SGPs.

In our study, we also revealed a novel level of LTR-RTE niche partitioning, at the integration step of the replication cycle. It is well-documented that different TE species, belonging to various classes, exhibit diverse target site preferences due to distinct transposition mechanisms^8,21,44^. For example, DNA transposons, like the P-element, manage to create new copies by integrating near the replication origins of the Drosophila genome^9,21^, whereas retrotransposons Ty1 and Ty3 specifically insert into Pol III promoters of *S. cerevisiae*^8^. Recently, it has been suggested that LTR-RTEs are rather attracted to open chromatin of active genes, whereas LINE elements, such as the I-element, target AT-rich sites and tend to integrate near telomeres^9^. These TE-specific host affinities have been described to depend on the enzymes driving their integration such as transposases or integrases. We found here that even LTR-RTEs of the same class, despite using the same integration mechanism, preferentially integrate into open chromatin domains harboring distinct chromatin features (Figure 3). This finding suggests that each LTR-RTE species has evolved specific interactions between its integrase and host co-factors (DNA- and/or chromatin-binding proteins) providing different affinities for specific genetic and epigenetic marks.

Note that, although specific for each LTR-RTE, their preferred epigenetic landscapes share a common feature, open chromatin, a permissive location for subsequent efficient transcription. These similarities might be considered as cases of concerted evolution by sharing general molecular mechanisms of targeting. A famous example of decompacted chromatin targeting is the histone H4 tail that can no longer be targeted by HIV when embedded in closed chromatin^45,46^.

Our data suggest that the specific integration of LTR-RTEs into distinct epigenetically defined domains might, at least partly, result from different integration timings during development. Strikingly, gtwin and ZAM landing site landscapes correlated with chromatin accessibility data sets extracted from early and late stages of embryogenesis, respectively. The hypothesis, assuming replication cycles with different timings of integration, is supported by the second wave of ZAM expression observed later in embryonic gonads. An obvious difference between these two putative cellular integration niches concerns their ability to proliferate. It is worth noting that, unlike early embryonic nuclei that are rapidly cycling, gonadic germ cells are no longer dividing. As it has been suggested for HIV, further experiments will be necessary to know whether gtwin and ZAM integrases have distinct abilities to be imported into non dividing nuclei.

### Putative selective forces leading to niche partitioning

By studying the simultaneous replication of four LTR-RTEs in the Drosophila germline, we observed distinct patterns suggesting that these LTR-RTEs occupy different ecological niches within the TE community. Here, we briefly speculate about the putative selective forces that might have led to niche partitioning.

First, we can notice that the expression of the four LTR-RTEs species appears to be restricted to gonadal tissues, the only host compartment supporting vertical transmission of the new TE copies. On the contrary, replication in non-gonadal tissues is not only useless for the TE replication but could have been counter-selected by the host as a possible cause of diseases like cancer and aging-related decline^47,48^. A second type of selective pressure might have prevented toxic TE expression^48^ in the germline stem cells, the immortal cell lineage of the gonad. That is probably why ZAM and gtwin are expressed in differentiated somatic gonadal cells, while roo, despite being a germline-specific TE, is expressed in nurse cells, which are differentiated germ cells destined to disappear at the end of oogenesis. Third, on one hand, the new TE copies need to insert into the germinal genome, but, on the other hand, the resulting DNA damage may be even more deleterious for the germline survival than the toxicity of the expression step. As a possible trade-off, integration is delayed until the DNA damage-tolerant embryonic stage of development^49^ that is followed by larval stages where germ cell division may compensate for previous cell death^50^. Fourth, further research is needed to characterize the phenotypic effects of the TE insertions we studied and determine if their preferred integration sites correspond to TE-specific safe havens within the host genome. Similarly, it is unknown whether the TE-specificity of these integration niches results from detrimental fitness effects of competition between different TE species at the insertion sites. Finally, our non-overlapping TE expression patterns are in agreement with previous observations^7^ suggesting that such a competition between somatic TEs might have led to expression niche partitioning.. In conclusion, TE niche partitioning highlights the complex interplay of positive and negative selection forces applied to TEs and their hosts and leading to their stable coexistence.

## Materials and Methods

### Drosophila stocks

Fly stocks (G0, G11, G31, G73), used to determine LTR-RTE mobilization and integration, shared the same genotype: *w ; tj-GAL4 ; tubP-GAL80ts*, *sh-piwi*, as previously described^10^. These stocks have been initially shifted, at every generation, from 20°C to 25°C during a 5-day period, at the adult stage (Figure 1A). At the 11^th^ (G11), 31^th^ (G31) and 73^rd^ (G73) generation, a large subset of the shifted population was isolated to maintain a stock constantly at 20°C, the non-permissive temperature for piwi-sKD. A fraction of the initial parental line G0 was also kept constantly at 20°C for 100 generations (G0F100). At each generation, the strains were maintained with a large progenitor population of more than 500 flies. A Drosophila line harboring the genotype: *w* ; *vas::EGFP* was used^39^.

### Analysis of female fecundity

For the control (Charolle) and the tested (G0F100) strain, 20 freshly hatched females were mated with 10 males at 20°C for 3 days while 10 freshly hatched females were mated with 5 males for 4 days at 25°C. The flies were then let to lay at 20°C and 25°C, respectively, and the number of F1 eggs was counted every 24 hours for 3 days. All egg collections were then left to develop at 20°C. The number of F1 pupae was counted for each condition. The number of F2 eggs laid by 20 3-day-old F1 mated females was counted at 20°C for both conditions. Three biological replicates were analyzed.

### Oxford Nanopore Technology (ONT) Sequencing Data Analysis

DNA was extracted from 100 males as previously described^51^ and long-read sequencing data were analyzed using the TrEMOLO software^11^ with some modifications. To detect newly integrated transposable elements, we employed the OUTSIDER TE detection module with, as a reference, the Dmel_R6.32 reference genome from FlyBase (v.104). Settings parameters for size and identity were set at 80%. The LTR-RTE database was extracted from the collection of reference TEs from Bergman’s laboratory (https://github.com/bergmanlab/transposons). The quality of the reads was analyzed in Table S5. Frequency estimation was conducted using the TE analysis module of TrEMOLO and reads identified as clipped reads by TrEMOLO were excluded from the frequency calculation.

### Annotation of false positive new insertions

The G0F100 library and the shifted libraries were established from populations that independently evolved from a shared G0 ancestor line. Consequently, any insertions found in both the G0F100 and any shifted library were attributed to the G0 parental genome. This allowed us to annotate as false negative pre-existing insertions those that were likely missed in the low quality G0 parental library, characterized by low coverage and shorter reads. All annotations were performed on the Dmel_R6.32 reference genome from FlyBase (v.104).

### Annotation of newly integrated LTR-RTE in piRNA clusters

The piRNA clusters were annotated on the Dmel_R6.32 reference genome using the published database https://www.smallrnagroup.uni-mainz.de/piRNAclusterDB/data/FASTA/Drosophila_melanogaster.piRNAclusters.gtf). Then a comparison between piRNA cluster coordinates and the LTR-RTE coordinates was used to determine the presence of new insertion in piRNA clusters.

### Single-molecule inexpensive RNA fluorescence *in situ* hybridization (smiRNA-FISH) probe preparation

39-48 probes of 20 nucleotides targeting specifically ZAM, gtwin, roo or copia transcripts were designed using Oligostan script^52^. Primary probes were produced in 96-well plates. For convenience, the oligonucleotides are delivered in Tris-EDTA pH 8.0 (TE) buffer, at final concentration of 100μM. An equimolar mixture of the different primary probes was prepared and diluted 5 times in TE buffer to obtain a final concentration of 0.833μM for each individual probe. Fluorescent labeled FLAP-X (5’-Cy3/CACT GAG TCC AGC TCG AAA CTT AGG AGG/Cy3-3’ or FLAP-Y (5′-Cy3/AA TGC ATG TCG ACG AGG TCC GAG TGT AA/Cy3-3′) were delivered lyophilized and resuspended in TE buffer at final concentration of 100μM. The reverse complement of each of these respective sequences was added at the 3’end of each specific probe (Table S6). Annealing between specific probes and their respective FLAP was performed as previously described^52^ and then diluted in hybridization buffer.

### smiFISH in ovaries and embryos

Ovaries were dissected in PBS1X and fixed during 20 minutes in PBS-Triton 0,3% (PBS-Tr) containing 4% formaldehyde. After several washes in PBS-Tr, ovaries were immersed in 100% methanol by successive baths in PBS-Triton 0,3% solution containing an increasing percentage of methanol. At this stage, ovaries can be kept in methanol at −20°C for several weeks. Embryos were collected and dechorionated in 2.6% bleach. They were rinsed extensively with water and fixed in 1:1 volume of fixative solution (4%Formaldehyde, KCl 60mM, Nacl 150mM, spermidine 0,5mM, Spermine 0,15mM, EDTA 2mM, EGTA 0,5mM, PIPES 15mM) and heptane for 25min at room temperature with agitation. Upon removal of the aqueous phase, an equal volume of 100% methanol was added before a vortexing for 1 min. Devitellinized embryos were collected from the methanol phase and then washed 3 times with 100% methanol. At this stage, embryos can be kept in methanol at −20°C for several weeks. Fixed embryos or ovaries were first washed twice in 50% methanol/50% ethanol for 5 minutes, rinsed twice in 100% ethanol and then washed two times in 100% ethanol for 5 minutes. They were incubated in blocking buffer (PBS 1X, tween 0,1%, RNAsin, BSA 0,2mg/mL in nuclease-free H_2_O) for 1 hour (a wash every 15 minutes) and once in wash buffer (SSC 2X, deionized formamide 10%, H2O in nuclease-free H_2_O) before the O/N incubation at 37°C at 350 rpm with smiFISH probes (Table S6) and either an anti-Rat Vasa antibody (DHSB, 1:120) or a Guinea Pig traffic jam antibody (gift from D. Godt^53^, Toronto, 1: 120) diluted in the hybridization buffer (10% deionized formamide, 2X SSC, 100mg tRNA, 5% dextran sulfate, 2mM VRC (NEB), 0,2mg/mL BSA). Subsequently, embryos/ovaries were washed with a wash buffer twice for 1 hour at 37°C and once for 1 hour at room temperature. Embryos were transferred in PBS, 0,1% Tween (PBT), 10% donkey serum and either Donkey anti Rat Alexa 488 (Molecular probes) or Donkey anti Guinea Pig Alexa 594 (Molecular probes) was added at 1:500 dilution. After several washes in PBT and DAPI staining, embryos/ovaries were mounted in ProLong Gold Antifade mounting medium (Thermo Fisher Scientific). smiFISH coupled with vasa immunostaining was imaged with LSM 880 with Airyscan module (Zeiss) using 40X/1.4 N.A objective. Airyscan processing was performed using 2D Zen Black v3.2 (Zeiss) prior to analysis. smiFISH coupled with traffic jam immunostaining was imaged with Leica SP8 confocal microscope equipped with 40X/1.4 N.A objectives. Image acquisition was done with the following settings: 2048x2048 pixels or 1024x1024 pixels, 16-bit depth.

### Distribution of LTR-RTEs insertions relative to genomic structures and chromatin states

Using the chromosomal gene and exons annotations of *Drosophila melanogaster* genome (BDGP6.46) available on Ensembl Biomart^54^ except for the Y chromosome, we partitioned the genome in three mutually exclusive regions corresponding to exons, introns and intergenic regions. Exons were already annotated in a bed file^54^. Introns were defined as genomic regions that are present in the gene bed file and which are not in the exon bed file. Intergenic regions are defined as genomic regions that do not overlap with the gene bed file. Using this partition and our annotations of LTR-RTEs insertion sites, we then determined the number of copia, roo, gtwin and ZAM insertions occurring in these three categories of genomic regions (Table S4). To determine whether a specific structure (Intergenic, Intron and Exon) is enriched or depleted for insertions of each considered LTR-RTE, bilateral binomial statistical tests were performed. To do so, the size of each structure relative to the genome was computed using bedtools genomecov^55^ default parameters, defining the relative size of intergenic regions (pig=0.314359), introns (pin=0.418317) and exons (pex=0.267325) (Table S4). Null hypothesis corresponds to the probability for each LTR-RTE species to be inserted in each defined structure due to its proportion in the genome. We used the Benjamini-Hochberg step up procedure to control the false discovery rate (FDR), which is defined as the expected value of the proportion of erroneous rejection of the null hypothesis when conducting multiple comparisons.

Chromatin state annotations previously published^18^ based on dm3 genome version were transformed to the latest version (Dmel_R6.32) using liftOver tool^56^. As for genomic structures, the proportion of each chromatin state relative to the genome was computed using bedtools genomecov^55^ default parameters (Table S4). Significant enrichment or depletion of LTR-RTE insertions in the different chromatin states were calculated using bilateral binomial statistical test considering the null hypothesis is the probability of insertion in a given state for each LTR-RTE species is equal to the relative size of this state within the genome. As in the previous part, we used the Benjamini-Hochberg step up procedure to control the false discovery rate (FDR).

### Relationship between the orientation of LTR-RTE and gene expression direction

The significance of the LTR-RTEs orientation within genes, according to gene annotation of the Drosophila genome (BDGP6.46) available on Ensembl Biomart^54^, was evaluated using binomial tests corrected by Benjamini–Hochberg with equal probabilities for insertions to occur in either orientation.

### Analysis of ATAC-seq and ChIP-seq available datasets

Raw data from published ATAC-seq and ChIP-seq experiments (Table S7) were analyzed to generate BigWig files using deepTools (v3.5.4.post1) bamcoverage package with default parameters, excluding regions (-bl) identified as blacklisted in dm6^57^. BigWig files were first used with the ComputeMatrix package with reference-point mode from deepTools to filter and sort regions based on their scores in order to compute signal distributions centered on the LTR-RTE insertion sites in a region spanning 2kb upstream and downstream of the insertion. The mean number of reads across the 4kb window was calculated using the (--averageTypeBins) option from ComputeMatrix, with a 50bp interval. On the other hand, BigWig files were converted into BedGraph format using the UCSC tool bigWigToBedGraph^58^. macs2 (v2.2.7.1) (bdgpeakcall package^59^ was used to perform a peak calling to generate bedfiles with the following parameters: (--cutoff) manually set; (--min-length) 60 and (--max-gap) 150. For the transcription factors ChIP-seq (Table S7), a consensus bed file was created by keeping only overlapping regions from the different replicates using bedtools intersect (v2.27.1)^55^. Random profile was generated using 100 random profiles each corresponding to an average profile obtained from 100 random positions (4kb window). We used ComputeMatrix with reference-point mode from deepTools as described above. PlotProfile with the (--outFileNameData) option was used to obtain each distribution of the average read number for the 100*100 randomly selected positions generated bed files. Finally, the mean number of read matrix was computed and used with deepTools plotProfile for visualization.

### Statistical analysis of available sci-ATAC-seq datasets

BigWig files from the sci-ATAC-seq atlas previously published^36^ were analyzed based on two criteria: they must represent an identified cell type and cover at least 70% of genomic data. ComputeMatrix package was used to assess the average chromatin accessibility around the 210 insertions of gtwin and the 101 insertions of ZAM as previously described. To determine the number of gtwin insertions shared between the first four clusters found in the 0-2 hour window of embryonic development, bed files were created as described before and visualized with a Venn Diagram.

To analyze globally chromatin accessibility throughout the 8 time windows of embryonic development, ATAC-seq signals (200 bp window) centered around ZAM or gtwin insertions were averaged for each defined cluster of each time window. The same technique was applied to 100 randomly selected regions of a 200 bp window. Ratios between the averaged ATAC-seq ZAM (or gtwin) signals and random ones result in a single data per cluster in a defined time window. Data corresponding to the same time window were used to generate boxplots and statistical analysis.

### Isolation of embryonic cells and cell sorting by flow cytometry

The embryonic cells were isolated as previously described^40^. Briefly, overnight laid embryos from *vas*::EGFP line^39^ were collected at 25°C and dechorionated in 2.6% bleach. Dechorionated embryos (i.e 400 mg) were transferred in a 7 mL Tenbroeck tissue grinder WHEATON™ filled with 6 mL of Schneider’s insect medium for homogenization with 2 slow strokes before a 700g centrifugation for 10 minutes at 4°C. The pellet was resuspended in 4 mL of PBS 1X containing 0.1% of Trypsin-EDTA and incubated at room temperature for 20 minutes. The addition of 4 mL of ice-cold PBS 1X containing 20% fetal bovine serum is sufficient to stop Trypsin reaction before a 700g centrifugation for 10 minutes at 4°C. Pellet containing separated embryonic cells was resuspended in Schneider’s insect medium (2mL) and filtered in a 40µm mesh before the addition of 1 mL of Schneider’s insect medium. A final filtration in a 20µm mesh was performed before cell sorting by flow cytometry. Embryonic cellular samples were analyzed using a 4-Laser-V16-B14-R8YG10 Aurora spectral cell sorter (Cytek, Biosciences, USA) to sort GFP-positive Primordial germ cells (PGC) from GFP negative somatic cells through a measurement of complete fluorescence spectrum of individual cells. GFP signal was determined by a 488 nm excitation line and detected in its full spectrum emission with B1 as peak channel (498nm-518nm). 2.5 x 10^5^ events were recorded per sample and analyzed using the SpectroFlo software version 1.2.1 (Cytek, Biosciences USA). To define and sort the target cell populations (GFP-positive cells), three sucessive steps of gating were applied. First, cells were gated using the two physical parameters FSC and SSC excluding dead cells and debris. Second, doublets were excluded by comparing the width versus the area of SSC and FSC. Finally, FSC dot plot and GFP signal reported as percentage in positive or negative cells were used to gate and sort the two populations. Live cell sorting experiments were performed at 4°C with a 70μm nozzle that allows sorting at high speed (2 x 10^4^ events per second). Sorted cells were collected into PBS containing 20% of Fetal Bovine serum (FBS) prior to a final centrifugation at 700g at 4°C and a −80°C freezing in DMSO supplemented with FBS.

### ATAC-Seq experiments

ATAC-Seq experiments were performed using the ATAC-Seq kit from Diagenode (catalogue no. C01080002). Input material was between 100,000 to 130,000 cryopreserved PGCs (GFP-positive) cells isolated from whole embryos. Tagmentated DNA was amplified by PCR using 13 cycles and the purified DNA libraries were sequenced (paired-end sequencing 150 bp, roughly 2 Gb per sample) by Novogene (https://en.novogene.com/). ATAC-Seq were performed in duplicates, following Encode’s standards (https://www.encodeproject.org/atac-seq/#standards).

### ATAC-Seq analysis

After initial quality checks of the data using FastQC (v0.12.1), the adapters (CTGTCTCTTATACACATCTNNNNNNNN) were trimmed using cutadapt (v4.2). Cleared reads were aligned to the *Drosophila* genome (Dmel_R6.32 release) using bowtie2 (v2.5.1). Duplicates alignements were removed using the fixmate and markdup packages of samtools (v1.17). The read coverage normalized by RPKM (--normalizeUsing) were computed using bamCoverage package from deepTools (v3.5.4.post1) with default parameters, excluding regions (-bl) identified as blacklisted in dm6. The chromatin accessibility in a region spanning 1kb upstream and downstream of ZAM insertions and 100 random genomic positions were computed with the ComputeMatrix package from deepTools as described above. The mean signal obtained from the duplicate were then computed using Matlab (R2024).

### Statistical analyses and visualization

Statistics and data visualization were performed using the ggplot2 library (v3.4.3) (https://ggplot2.tidyverse.org) on R (v4.3.1) (https://www.R-project.org/) and Matlab (R2024). Cytoscape (v3.10.1)^60^ was used to create a graph.

## Data availability

Long reads sequencing data previously published and presented in this study have been deposited at ENA (https://www.ebi.ac.uk/ena) under the accession numbers ERP122844 and PRJEB75331 respectively. The source code of TrEMOLO as well as all the accessory codes are available at https://github.com/Drosophila GenomeEvolution/TrEMOLO. The ATAC-seq raw data are available on GEO under accession number: GSE274394.

## Acknowledgments

We thank Bernd Schuettengruber for the comments and the editing of the manuscript, Callum Burnard and Gonzalo Sabaris for scientific discussions. We acknowledge the ISO 9001 certified IRD itrop HPC (member of the South Green Platform) at IRD Montpellier for providing HPC resources that have contributed to the research results reported in this paper (URLs: https://bioinfo.ird.fr/ and http://www.southgreen.fr); the Genotoul platform (https://genotoul.fr/) and (https://www.france-bioinformatique.fr/) for providing calculation time on their servers; BioCampus MRI platform for microscopy and Drosophila core facilities. We thank Akira Nakamura for the drosophila line *w* ; *vas::EGFP*, Makoto Hayashi for his help on the PGC isolation protocol, Felicia Leccia from the MRI-Cyto IRMB Cytometry platform for the cell sorting and Bernd Schuettengruber for his help on the ATAC-seq experiments.

## Author contributions

Conception: S.C; A.P.; Computational analysis of NGS and genomics data, M.M.; M.V.; Statistics, D.G.; M.V.; Experiments C.G, M.L., B.M., M.V.; Methodology and analyses: B.M.; C.G.; A.P.; S.C.; Supervision, C.G.; S.C.; Visualization: B.M.; D.G.; M.V.; Writing: C.G.; A.P.; S.C.; Funding & infrastructure: S.C.

## Supplemental figures

**Figure S1: Female fertility at 25°C and 20°C**

The abundance and fecundity (left: F1 eggs; middle: F1 adults; right: F2 eggs) of the progenies of G0F100 (blue) and Charolle (ctrl, light blue) flies shifted at 25°C were normalized with those of the same females kept at 20°C. F1flies were allowed to develop and lay at 20°C. This experiment was technically repeated three times using three independent biological replicates at 25°C and 20°C, except for F2 eggs laid where only one technical replicate per biological replicates was considered. Each bar corresponds to the mean value and errors bars correspond to standard deviation.

**Figure S2: New LTR-RTE insertions are not positively selected in G73 generation**

(A) Metaplots depicting the frequency of each new LTR-RTE insertion according to its positioning along each chromosome. Chromosomes 2 and 3 are separated in two metaplots corresponding the left and right arms of these chromosomes. Each family of LTR-RTEs is represented with a color code indicated on the top right-hand side. (B) Table indicating the numbers of LTR-RTE insertions in the different piRNA clusters. (C) Frequency of each LTR-RTE insertion found in piRNA clusters.

